# Ginsenoside Rg1 defenses PC-12 cells against hydrogen peroxide-caused damage via up-regulation of miR-216a-5p

**DOI:** 10.1101/662817

**Authors:** Guangkun Yi, Li Liu, Chaoliang Lv, Yanchun Wei, Tingzhen Yan

## Abstract

**Background:** Spinal cord injury (SCI) is a destructive trauma accompanying with local injury, half of which cause chronic paralysis. Ginsenoside Rg1 exerts anti-apoptosis and anti-autophagy properties. Therefore, our goal was to study the protective mechanism of Rg1 in attenuating cell injury.

**Methods:** MiR-216a-5p inhibitor was transfected into PC-12 cells, then cells were pre-treated by Rg1 and treated with 300 μM hydrogen peroxide (H_2_O_2_) for 24 h. CCK-8 and apoptosis experiments were done to test cell activity and apoptosis respectively. Expression of miR-216a-5p and cell damage relative factors was tested via qRT-PCR and western blot experiments, respectively.

**Results:** H_2_O_2_ induced cell activity suppression, apoptosis and autophagy well at the concentration of 300 μM, leading cell injury. Rg1 could attenuate cell injury induced by H_2_O_2_ at the working concentration of 200 μM that it elevated cell activity, attenuated apoptosis and autophagy and activated PI3K/AKT and AMPK signal pathways. Further, miR-216a-5p was up-regulated by Rg1. Rg1 played its role in relieving cell injury by positively regulating miR-216a-5p.

**Conclusion:** Our study demonstrated that Rg1 attenuated H_2_O_2_-caused cell injury through positively regulated miR-216a-5p.

## Introduction

Spinal cord injury (SCI), a common and destructive trauma (Hyun and Kim, 2010), is mainly caused by external forces such as lateral bending, excessive stretching, rotation, axial load and excessive bending, resulting in motor dysfunction, paralysis and other symptoms (van den Berg et al., 2010). Because of the limited therapy selection, the administration and care of SCI patients places a heavy burden on patients and caregivers. Of particular note, more than 60% of damages occur at the cervical level (Cripps et al., 2011), and the lifetime care costs are about at $1.1-$4.2 million per patient (Krueger et al., 2013). So, the precaution, curing and recovery of SCI are a major topic in the medical area. SCI relates two different stages of tissue injury, called primary and secondary hurt (Badner et al., 2019). Local tissue injury is caused by SCI and is important in secondary hurt in SCI (Fu et al., 2018), leading to apoptosis with loss of neurological roles. So, mechanism study of local injury after SCI is highly significant for curing SCI.

Ginsenosides, considering as one of the main pharmacological active ingredients of ginseng, is a steroid compound (Xiang et al., 2008). Ginsenoside contains the Panaxatriol (Rg1, Rg2, Re and Rf) and Panaxadiol (Rb1, Rb2, Rc and Rd) classes (Zhang et al., 2012a). Many beneficial effects of Rg1 have been proved in disorders such as hypertension (Chen et al., 2012), hypoxia/reoxygenation (Zhang et al., 2012b), Alzheimer’s disease (Huang et al., 2012) etc. Importantly, it has been reported that Rg1 exerts roles in inhibiting cell apoptosis, thereby exhibiting notable cardioprotective effects against I/R damage through a variety of mechanisms (Lee and Kim, 2014). Besides, Rg1 counteracts the aging of endothelial progenitor cells (Shi et al., 2011) and human fibroblasts (Zhou et al., 2012) and exerts a notable influence in suppressing cardiomyocytes and renal tubular cells’ autophagy (Mao et al., 2016). The influence of Rg1 in local injury after SCI still has been unknown yet.

MicroRNAs (miRNAs), short (22 nucleotides in length) non-coding RNAs, involve in many biological processes (Jiang and Chen, 2012), such as differentiation of ordinary tissues and are important in the pathogenesis of lots of human cancers (Taucher et al., 2016). MiR-216a-5p, known as an oncogene, involved in the progression of many cancer subtypes (Chen et al., 2018). Chen *et al.* has proved that miR-216a-5p elevates cell proliferation, activity and motility, and inhibits apoptosis (Chen et al., 2018). This finding demonstrates that miR-216a-5p has a positive effect on cell viability and anti-apoptosis. So it could be interesting to investigate if exerts regulation relation of miR-216a-5p and Rg1 in cell injury after SCI. Based on the above questions and guesses, we probed mechanism of Rg1 against H_2_O_2_-caused cell damage in PC-12 cells.

## Materials and Methods

### Cell

PC-12 cells were bought form Kunming Institute of Zoology (Kunming, China) in this whole study. Seed cells at a denseness of 1 × 10^4^ cells/ml in Dulbecco’s Modied Eagle Medium (DMEM)/F-12 medium (Gibco, Carlsbad, CA, USA) adding with 10% fetal bovine serum (FBS, Gibco), 100 μg/ml streptomycin and 100 U/ml penicillin (Gibco). Cells were kept in a wet incubator carried 5% CO_2_ and 95% air at 37°C. Change fresh medium every day. Ginsenoside Rg1 (analysis level of 97% pureness) was bought from Sigma-Aldrich (St. Louis, MO, USA), solubled in ethanol and stored at −20°C. Pre-treatment of cells with Rg1 for 1 h, and then were treated with a series of consistences of hydrogen peroxide (H_2_O_2_) for 24 h.

### CCK-8 experiment

A Cell Counting Kit-8 (CCK-8, Dojindo Molecular Technologies, Gaithersburg, MD, USA) was to test cell activity. Seed cells in 96-well plate with 5000 cells/well, and then add CCK-8 solution, keep cells in a wet environment carried 95% air and 5% CO_2_ for 1 h at 37°C. Absorbance was tested at 450 nm via a Microplate Reader (Bio-Rad, Hercules, CA, USA).

### Apoptosis experiment

Apoptosis analysis was done through propidium iodide (PI) and fluorescein isothiocynate (FITC)-conjugated Annexin V staining (BD Pharmingen, San Diego, CA, USA). Cells were cleaned in phosphatebuffered saline (PBS) for three times and stained in PI/FITC-Annexin V with 50 μg/ml RNase A (Sigma-Aldrich). Keep cells in dark processing at the room temperature for 1 h. Flow cytometry analysis was made through FACS can (Beckman Coulter, Fullerton, CA, USA). Data was analyzed via FlowJo software (Tree Star Software, San Carlos, California, USA).

### Transfection

MiR-216a-5p inhibitor and its relative NC were compounded by Life Technologies Corporation (Carlsbad, CA, USA) and transferred into cells. Transfection was done following the Lipofectamine 3000 reagent (Life Technologies Corporation). 48 h post-transfection was regarded as harvest moment in following assays.

### qRT-PCR

Overall RNA was extracted through Trizol reagent (Life Technologies Corporation) and handled with DNaseI (Promega, Madison, WI, USA). Taqman MicroRNA Reverse Transcription Kit and Taqman Universal Master Mix II with the TaqMan MicroRNA Assay (Applied Biosystems, Foster City, CA, USA) were to test miR-216a-5p expression. U6 was taken as inside comparison.

### Western Blot

Overall protein was extracted through RIPA lysis buffer (Beyotime Biotechnology, Shanghai, China) with protease inhibitors (Roche, Basel, Switzerland), and then quantified through BCA™ Protein Assay Kit (Pierce, Appleton, WI, USA). A Bio-Rad Bis-Tris Gel system was taken to build up a western blot system. Primary antibodies specific against Bax (ab32503, Abcam, Cambridge, MA, USA), pro-caspase-3 (ab183179), cleaved-caspase-3 (ab49822), pro-PARP (ab32064), cleaved-PARP (ab4830), β-actin (ab8226), beclin-1 (ab62557), p62 (ab56416), LC3-I and LC3-II (ab48394), t-PI3K (ab140307), p-PI3K (ab182651), t-AKT (ab179463), p-AKT (ab38499), t-AMPK (ab131512) and p-AMPK (ab23875) were readied in 5% blocking buffer. Primary antibody was cultured with membrane at 4°C all the night, then washing and incubating with secondary antibody, marking by horseradish peroxidase for 1 h at room temperature. Then the Polyvinylidene Difluoride (PVDF) membrane taken along blots and antibodies were transferred into the Bio-Rad ChemiDoc™ XRS system, adding 200 μl Immobilon Western Chemiluminescent HRP Substrate (Millipore, MA, USA) to shroud film surface. At last, semaphores were seized and strength of strip was quantified via Image Lab™ Software (Bio-Rad).

### Statistical analysis

All assays were duplicated for 3 times. Our consequences of multiplex assays are revealed as mean ± SD. Statistical analysis was done via Graphpad Prism 6.0 (GraphPad Software Inc., La Jolla, CA, USA). *P*-values were counted via a one-way analysis of variance (ANOVA). *P*-value of < 0.05 indicated statistical significant data.

## Results

### Rg1 extenuated H_2_O_2_-induced cell activity suppression and cell apoptosis

PC-12 cells were treated in various H_2_O_2_ consistences. From **Figure 1A**, we found that H_2_O_2_ had notably inhibiting effect on cell viability when the concentration was 100 (*P* < 0.05), 200 (*P* < 0.05), 300 (*P* < 0.01), 400 (*P* < 0.01) and 500 μM (*P* < 0.001). We chose 300 μM as the working concentration in the later assays because this was cell viability semi-lethal concentration. Besides, we tested effect of H_2_O_2_ on cell apoptosis. We found that apoptosis was notably increased by H_2_O_2_ (*P* < 0.001, **Figure 1B**). Similarly, apoptosis relative factors (Bax, cleaved-caspase-3 and cleaved-caspase-PARP) were obviously enhanced through H_2_O_2_ (**Figure 1C**), and standards of these factors were notably raised (all *P* < 0.01, **Figure 1D**). We got that H_2_O_2_ caused cell activity suppression and apoptosis.

**Figure 1.**
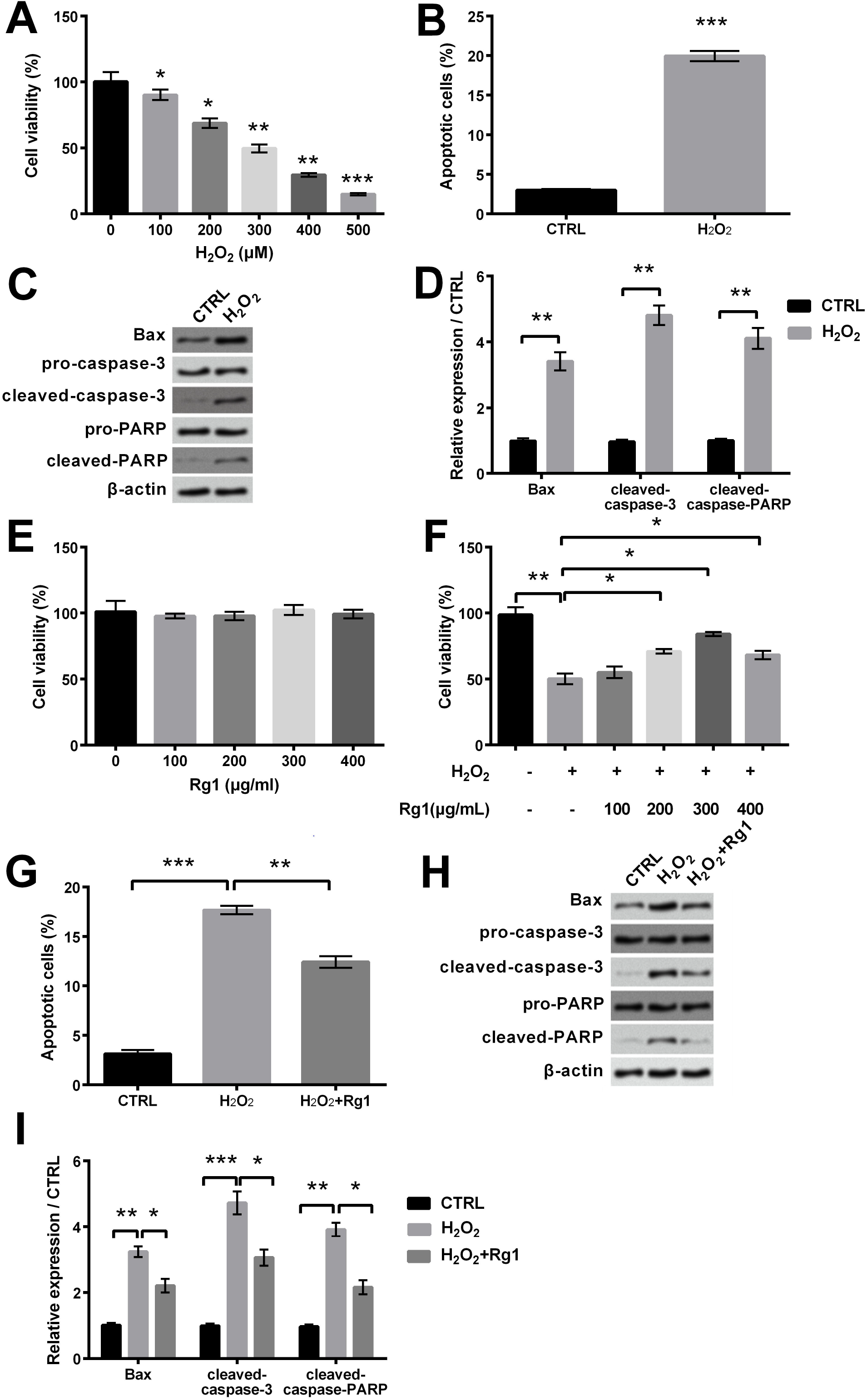
Influence of Rg1 in cell activity and apoptosiscaused by H_2_O_2_ in PC-12 cell. (A) Cell viability was tested under diverse consistences of H_2_O_2_ (0, 100, 200, 300, 400 and 500 μM). 300 μM was chose in the following experiments. (B) Cell apoptosis treated by H_2_O_2_ was tested via flow cytometry. (C) Apoptosis relative elements expression was tested via western blot. (D) Level of apoptosis relative factors was tested via western blot quantitative. (E) Cell viability was tested via CCK-8 by Rg1. (F) Rg1 attenuated H_2_O_2_-induced suppression of cell viability. (G) Rg1 attenuated H_2_O_2_-induced cell apoptosis. (H) Expression of apoptosis relative factors was tested via western blot. (I) Apoptosis relative elements standards were detected via western blot quantitative. * *P* < 0.05, ** *P* < 0.01 and *** *P* < 0.001 contrast with control and the indicated set.

For function of Rg1, following experimental results were clear. As shown in **Figure 1E**, there was no effect on cell viability by Rg1. We found that H_2_O_2_ could notably reduce cell viability (*P* < 0.01), whereas Rg1 could notably attenuate this reduction at 200, 300 and 400 μM (all *P* < 0.05). We chose 200 μM as the working concentration in the following experiments because this is the concentration when cell viability was half restored. Besides, for cell apoptosis, we found that Rg1 attenuated apoptosis induced by H_2_O_2_ (*P* < 0.01, **Figure 1G**). Similarly, **Figure 1H** revealed that expression of apoptosis relative factors was weakened by Rg1 compared with H_2_O_2_ group. Levels of these factors were raised through H_2_O_2_ (*P* < 0.01, *P* < 0.001 and *P* < 0.01), whereas Rg1 could decrease their levels (all *P* < 0.05, **Figure 1I**). So we got that Rg1 attenuated cell activity suppression and apoptosis induced by H_2_O_2_.

### Rg1 extenuated autophagy induced by H_2_O_2_

For autophagy, we tested three autophagy relative factors. Beclin-1 is autophagy gene and its overexpression can stimulate autophagy (Yue et al., 2003). Accumulation of p62 is a notable phenotype of autophagy-deficient tumor cells (Mathew et al., 2009). LC3-II is a marker for mature autophagosomes. Autophagy could be analyzed by testing the conversion of the autophagosome marker LC3-I to LC3-II (Wu et al., 2010). According to our results, **Figure 2A** showed the enhancement of beclin-1 and LC3-II/LC3-I through H_2_O_2_, while Rg1 could weaken this enhancement. Expression of p62 was weakened by H_2_O_2_, while Rg1 could eliminate this mitigation (**Figure 2A**). Besides, **Figure 2B** revealed the notable addition of beclin-1 and LC3-II/LC3-Ithrough H_2_O_2_ (*P* < 0.01 and *P* < 0.001), whereas were opposite by adding of Rg1 (*P* < 0.05 and *P* < 0.01). p62 expression was notably weakened through H_2_O_2_ (*P* < 0.05), whereas was increased by the adding of Rg1 (*P* < 0.05, **Figure 2B**). So we got that Rg1 could attenuate H_2_O_2_-induced autophagy.

**Figure 2.**
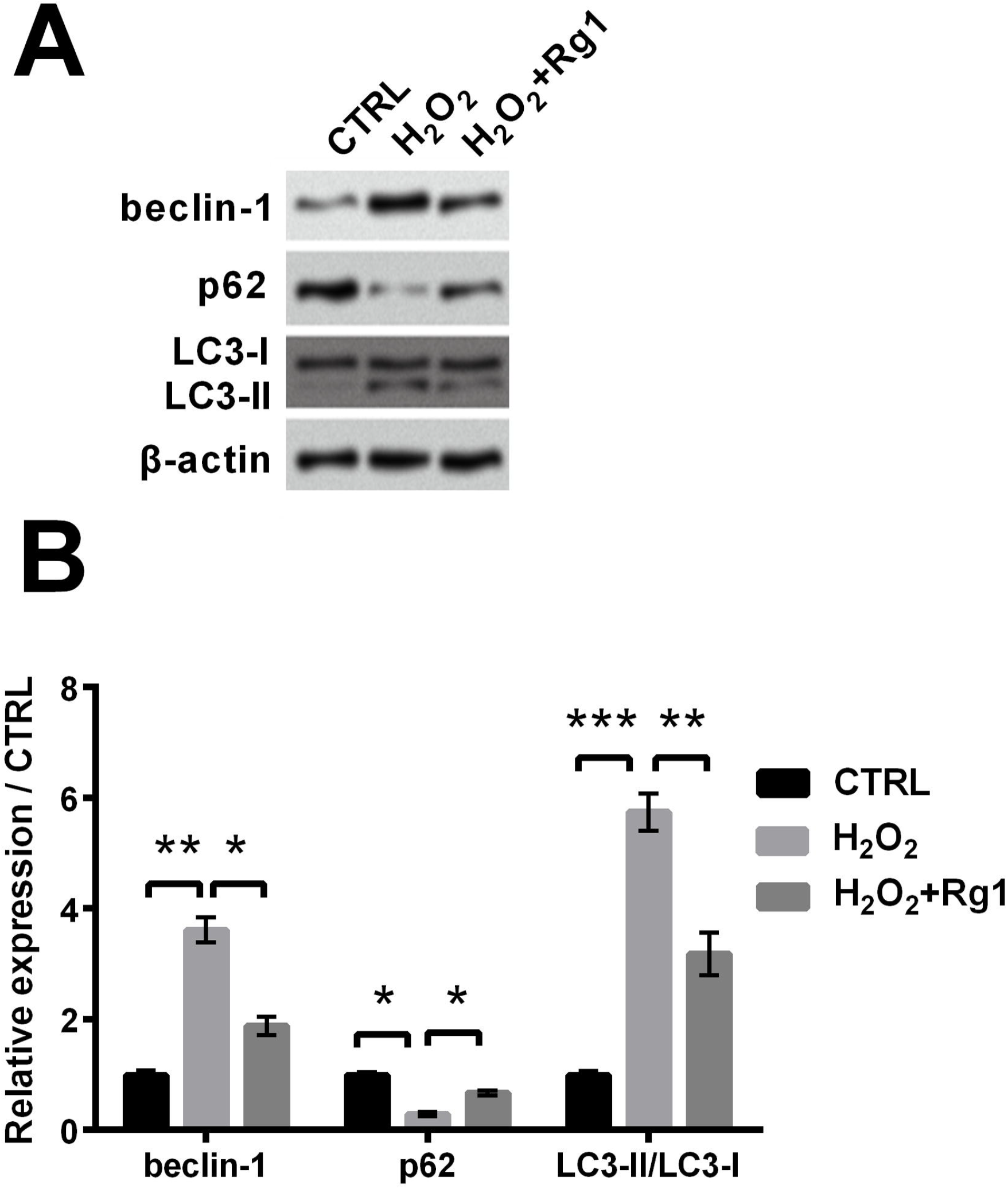
Influence of Rg1 in autophagy caused by H_2_O_2_. (A) Standards of autophagy relative factors were tested via western blot. (B) Standards of autophagy relative factors were tested via western blot quantitative. * *P* < 0.05, ** *P* < 0.01 and *** *P* < 0.001 contrast with indicated set.

### Rg1 positively regulated miR-216a-5p

From **Figure 3**, qRT-PCR assay indicated that miR-216a-5p was notably down-regulated after H_2_O_2_ treatment (*P* < 0.05). But, it was specifically up-regulated by adding Rg1 (*P* < 0.01). So we got that Rg1 up-regulated miR-216a-5p.

**Figure 3.**
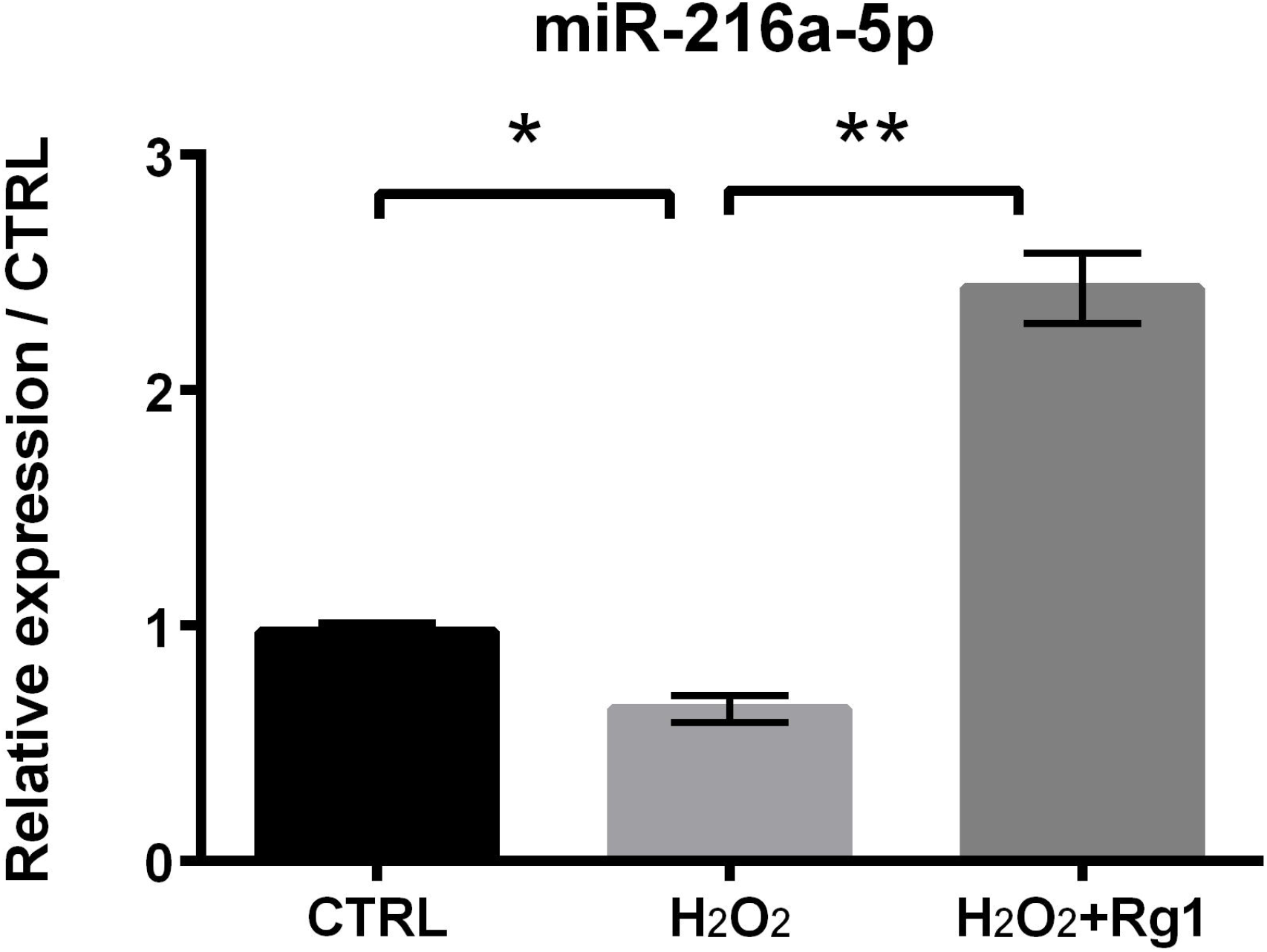
MiR-216a-5p was up-regulated by Rg1. mRNA standard of miR-216a-5p was tested via qRT-PCR. * *P* < 0.05 and ** *P* < 0.01 contrast with indicated set.

### Rg1 extenuated cell activity suppression, apoptosis and autophagy induced by H_2_O_2_ through up-regulating miR-216a-5p

qRT-PCR revealed that miR-216a-5p expression was notably suppressed after miR-216a-5p inhibitor transfection (*P* < 0.01, **Figure 4A**). Cell viability was notably aggravated by Rg1 contrast with H_2_O_2_ set (*P* < 0.05), whereas was notably alleviated when treated with Rg1 plus miR-216a-5p inhibitor (*P* < 0.05, **Figure 4B**). This result indicated that Rg1 attenuated H_2_O_2_-induced cell activity suppression by up-regulating miR-216a-5p. Besides, cell apoptosis was notably decreased by Rg1 contrast with H_2_O_2_ set (*P* < 0.01), whereas was notably increased when treated with Rg1 plus miR-216a-5p inhibitor (*P* < 0.01, **Figure 4C**). **Figure 4D-E** further indicated that levels of apoptosis relative factors were notably decreased through Rg1 contrast with H_2_O_2_ set (*P* < 0.05, *P* < 0.01 and *P* < 0.01), whereas were raised in Rg1-treated cells with miR-216a-5p inhibitor (*P* < 0.05, *P* < 0.01 and *P* < 0.05). So we got that Rg1 reduced H_2_O_2_-caused apoptosis by up-regulating miR-216a-5p. Additionally, **Figure 4F-G** revealed the notable attenuation of beclin-1 and LC3-II/LC3-I (*P* < 0.01 and *P* < 0.001) by Rg1 and level of p62 was increased (*P* < 0.05) by Rg1 constrast with H_2_O_2_ set. However, beclin-1 and LC3-II/LC3-I were notably enhanced (both *P* < 0.05) and level of p62 was notably decreased when treated with Rg1 plus miR-216a-5p inhibitor (*P* < 0.05). So we got that Rg1 attenuated H_2_O_2_-induced autophagy by up-regulating miR-216a-5p.

**Figure 4.**
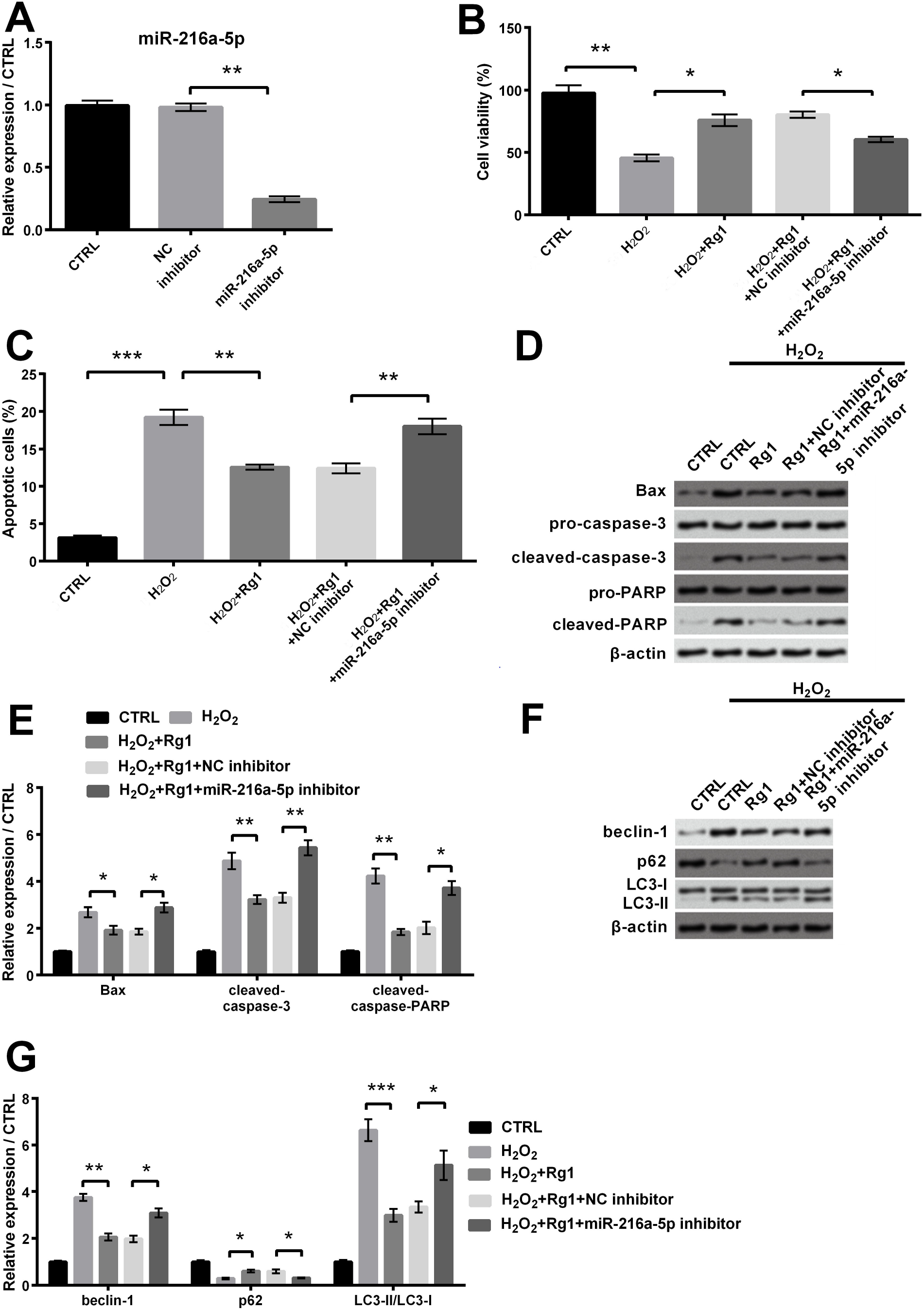
Protective effects of Rg1 via up-regulating miR-216a-5p after transfected with Rg1 plus miR-216a-5p and relative NC in PC-12 cells. (A) mRNA standard of miR-216a-5p was tested via qRT-PCR after miR-216a-5p inhibitor transfection. (B) Cell activity was tested via CCK-8. (C) Apoptosis was tested via flow cytometry. (D) Expression of apoptosis relative factors was tested via western blot. (E) Level of apoptosis relative factors was tested via western blot quantitative. (F) Standards of autophagy relative factors were tested via western blot. (G) Standards of autophagy relative factors were tested via western blot quantitative. * *P* < 0.05, ** *P* < 0.01 and *** *P* < 0.001 contrast with indicated set.

### Signal pathway

To further study the mechanism of Rg1, we focused on PI3K/AKT and AMPK signal pathways. **Figure 5A-B** indicated the notable addition levels of p-PI3K and p-AKT through Rg1 contrast with H_2_O_2_ set (*P* < 0.01 and *P* < 0.05), whereas were notably alleviated in Rg1-treated cells with miR-216a-5p inhibitor (*P* < 0.01 and *P* < 0.05). Besides, **Figure 5C-D** revealed that level of p-AMPK was notably aggravated by Rg1 compared with H_2_O_2_ group (*P* < 0.01), whereas was notably alleviated in Rg1-treated cells with miR-216a-5p inhibitor (*P* < 0.01). These results indicated that Rg1 elevated PI3K/AKT and AMPK pathways via positively modulating miR-216a-5p.

**Figure 5.**
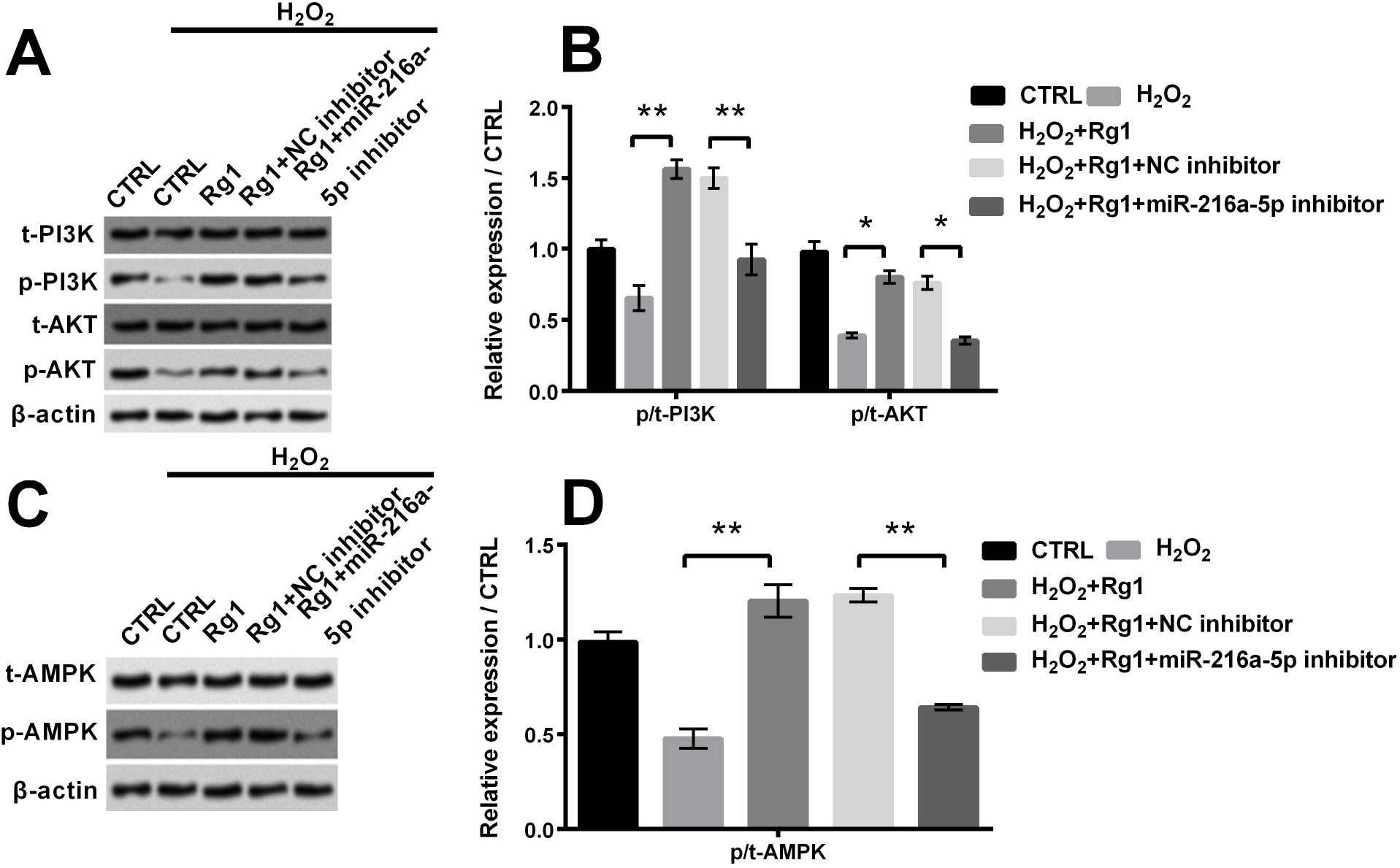
Rg1 elevated PI3K/AKT and AMPK signal pathways through positively regulating miR-216a-5p after transfected with Rg1 plus miR-216a-5p and relative NC in PC-12 cells. (A) Expression of PI3K/AKT pathway relative factors was tested via western blot. (B) Standards of relative proteins were tested via western blot quantitative. (C) Expression of AMPK pathway relative factors was tested via western blot. (D) Level of AMPK pathway related proteins were detected via western blot quantitative. * *P* < 0.05 and ** *P* < 0.01 contrast with indicated set.

## Discussion

SCI is a life-changing event. Recently, there is no effective treatment method to resume the functions of SCI patients. The complexity of SCI pathophysiology poses a huge challenge for researchers and clinicians seeking to develop therapeutic interventions (Bareyre, 2019). Local injury is the main event of secondary injury in SCI, eventually leading to apoptosis and ultimately loss of neurological function (Genovese et al., 2009). Therefore, defensing cells against local injury or relieving this injury can be an effective method to cure SCI. Rg1, an active component of ginsenosides, has been proved to exert positive effect on anti-apoptosis (Zu et al., 2016). Cell damage has been reported to be induced by H_2_O_2_ in cardiomyocytes (Zeng et al., 2019) and neural cells (Chen et al., 2016). Therefore, our study also found that H_2_O_2_ was a mediator capable of inducing damage in PC-12 cells, including activity suppression, promotion of autophagy and apoptosis. Our study firstly researched the attenuated mechanism of Rg1 in H_2_O_2_-caused damage in PC-12 cells. Rg1 was effective in defensing PC-12 cells against H_2_O_2_-caused damage. Rg1 could extenuate H_2_O_2_-induced cell activity suppression, apoptosis and autophagy, and active PI3K/AKT and AMPK pathways via positively regulating miR-216a-5p. Our results indicated that Rg1 may be an effective treatment of curing SCI.

Ginsenoside Rg1, the main bioactive ingredient in ginseng, has been proved to exert low toxicity that there was no change on cell viability and proliferation (Li et al., 2017). Much evidence indicates that Rg1 exerts beneficial effects, like anti-aging properties (Zhu et al., 2014). As we all know, Bax may control mitochondrial permeability transition and promote releasing cytochrome c, ultimately triggering the activation of caspases, leading to apoptosis (Li et al., 2017). Consistently, in our study, treatment of Rg1 reduced the level of Bax, at the same time, cleaved-caspase-3 and cleaved-caspase-PARP were weakened. These observations verified the anti-apoptosis function of Rg1. Additionally, autophagy is an important cellular process where cytoplasmic components are digested by lysosomes to keep cell homeostasis and energy production (Ravikumar et al., 2010). Rg1 has a notable pharmacological influence in suppressing autophagy (Mao et al., 2016). Our study were consistent with the report that Rg1 strongly inhibited autophagic factor (beclin-1 and LC3-II/LC3-I), and counteracted attenuation of p62 induced by H_2_O_2_, leading to inhibiting H_2_O_2_-induced autophagy in PC-12 cells. Consistently, our study indicated that Rg1 could counteract inhibition of cell activity induced by H_2_O_2_ and attenuate cell apoptosis and autophagy, suggesting the anti-oxidant and anti-autophagy functions of Rg1 in SCI.

To further study the mechanism of Rg1, we turn our attention to miRNA. MiRNA is important in cell growth, like proliferation and apoptosis (Ameres and Zamore, 2013). MiR-216a-5p, acknowledged as an oncogenic gene, is involved in tumorigenesis and development of human cancers (Liu et al., 2018). MiR-216a-5p significantly elevated cell activity and reduced apoptosis in H_2_O_2_-caused 16 HBE cells of Asthma, suggesting that miR-216a-5p could regulate H_2_O_2_-caused damage (Chaoyang et al., 2019). Besides, of interest, beclin-1 was the latent mark of miR-216-5p, which could inhibit ox-LDL-induced autophagy in human umbilical vein endothelial cells (HUVECs) through modulating levels of intracellular beclin-1 (Menghini et al., 2014). These reports indicate that miR-216a-5p not only elevates activity, suppresses apoptosis, but also inhibits autophagy, suggesting that miR-216a-5p may reduce H_2_O_2_-caused cell damage. Also, functions of miR-216a-5p are similar to those of Rg1. So it is worth to investigate if exerts a relation of Rg1 and miR-216a-5p. For the first time, our study found the regulation relationship between Rg1 and miR-216a-5p. We got that Rg1 could up-regulate miR-216a-5p to attenuate cell injury induced by H_2_O_2_. This finding is a major discovery in SCI research.

Furthermore, the biological process is inseparable from the regulation of signal pathways. It has been proved that AMPK/PI3K/AKT pathways are key coordinator protecting cells from oxidative and inflammatory damage (Lv et al., 2017). PI3K was reported to be related to many cellular functions, like proliferation and apoptosis (Rasul et al., 2012). AKT is a key downstream effector of PI3K and exhibits anti-apoptotic effects (Zheng et al., 2015). AMPK is present in metabolically related organs. Cellular metabolism stimulation like cell stress can active it (Zheng et al., 2015). Moreover, Lin *et al.* found that the formation of autophagosomes was accompanied by inhibition of the PI3K/AKT and AMPK signal pathways. This finding indicated that there was negative regulation between these two pathways and autophagy. Therefore, the above findings suggested that there might be positive effects of these two pathways on H_2_O_2_-caused cell damage. It is worth noting that our study verified this suppose. We firstly build up the relation among Rg1, miR-216a-5p and AMPK/PI3K/AKT pathways that Rg1 activated PI3K/AKT and AMPK signal pathways through positively modulating miR-216a-5p to reduce cell damage. This regulation mechanism provides a theoretical basis for attenuating cell injury after SCI.

## Conclusion

Our study firstly reported the underlying effects and mechanism of Rg1 in cell injury of SCI. We demonstrated that Rg1 could up-regulate miR-216a-5p, attenuate cell activity suppression, apoptosis and autophagy, and active PI3K/AKT and AMPK signal pathways to decrease H_2_O_2_-caused damage in PC-12 cells. Because local cell injury significantly aggravates SCI, we propose that Rg1, an effective biological macromolecular, may supply a novel therapeutic approach for curing SCI.

## Conflict of Interest statement

The authors declare that there are no conflicts of interest.

## Acknowledgments

None

## Fundings

This research did not receive any specific grant from funding agencies in the public, commercial, or not-for-profit sectors.

## Availability of data and materials

The dataset(s) supporting the conclusions of this article is(are) included within the article.

